# Multi-drug resistance bacteria predict mortality in blood stream infection in a tertiary setting in Tanzania

**DOI:** 10.1101/705863

**Authors:** Joel Manyahi, Upendo Kibwana, Edna Mgimba, Mtebe Majigo

## Abstract

**Background:** Blood-stream infections (BSI’s) are serious and life-threatening infections associated with high mortality and morbidity. In resource limited settings, there is paucity of data on predictors of outcome in patients with BSI. This study aimed at examining the predictors of mortality in patients with BSI as well as bacteria causing BSI.

**Methods and Materials:** This was cross-sectional study conducted in Muhimbili National Hospital between April and May 2018. Blood culture results from all inpatients at clinical microbiology laboratory were recorded and clinical information were retrieved retrospectively from the files. Bacteria from positive blood culture were identified and antimicrobial susceptibility was performed.

**Results:** The overall prevalence of BSI was 11.4% (46/402), with case fatality rate of 37%. There was significant high rate of BSI in patient who had died compared to those survived p= 0.008. Gram-negative bacteria (74%) were the common cause of BSI, with predominance of Enterobacteriaceae (22), followed by *Pseudomonas aeruginosa* (11). Majority (70.5%) of the bacteria isolated from patients with BSI were multi-drug resistant. Forty six percent of *Pseudomonas aeruginosa* were resistance to meropenem. Sixty eight percent 68.2% (15/22) of Enterobacteriaceae were ESBL producers. Carbapenemases production were detected in 27% (3/11) of *Pseudomonas aeruginosa* and in one *Proteus mirabillis*. Forty percent (40%) of *Staphylococcus aureus* were methicillin resistant *Staphylococcus aureus* (MRSA). Positive blood culture (cOR 7.4, 95%CI 1.24 – 43.83, p 0.03) and admission in ICU cOR 4 (95%CI 1.7 – 9.41, p 0.001) were independent factors for mortality in suspected BSI. Isolation of Multi-drug resistant bacteria was independent predictor for mortality in confirmed BSI (cOR 7.4, 95%CI 1.24 – 43.83, p 0.03).

**Conclusion:** The prevalence of BSI was 11.4%, with majority of bacteria in BSI were MDR. Positive blood culture and MDR were predictors for mortality.

## Introduction

Blood stream infection (BSI) is a life threatening and has been associated with increased mortality, morbidity and health care costs (1). Often, multi-drug resistant bacteria causing BSI attribute to disproportionate poor outcomes compared to susceptible bacteria (2, 3). The incidence and prevalence of BSI varies considerably between developed and developing countries (4–6). Nevertheless, epidemiology of BSI in both community and hospital settings is evolving. Besides that, data on rapid changing bacterial etiology of BSI in Tanzania is scarce.

Treatment of BSI in resource limited setting is largely empiric with broad spectrum antibiotics. Empiric treatments often fail to target the correct pathogens effectively, leading to treatment failures and increasing mortality (1, 7). To address these, clinical microbiology laboratories may play important roles on effective management of BSI. Prompt reporting of results coupled with identifying critical values and antibiogram pattern provided by laboratories facilitate on successful management of patients with BSI. Elsewhere, factors predicting mortality in BSI have been investigated (2, 8), but few data exist in Tanzania.

The increase burden of hospital acquired BSI caused by Multidrug resistant pathogens including extended spectrum beta-lactamases have been previously observed in Tanzania (3). However, in a world of evolving bacteria pathogens, updated data are warranted for improving management of BSI. Our study aimed at identifying the current bacterial etiology and predictors of mortality in BSI.

## Materials and methods

### Study design and setting

This was a cross-sectional study conducted at Muhimbili National Hospital (MNH) between April and May 2018. MNH is the largest tertiary hospital in Tanzania, serving approximately 6 million populations from Dar es Salaam. It has a 1500 bed capacity, attending approximately 1200 inpatients per week and approximately 1200 outpatients per day. MNH is also a training facility for Muhimbili University of Health and Allied Sciences and main referral hospital in Tanzania.

### Study Population

The study included all in-patients with clinical features suggestive of BSI, from whom blood specimens for culture were processed at MNH Clinical Microbiology Laboratory.

### Data collection

Structured data collection tool was used to record results of blood culture, colonial morphology, Gram stain, isolates identity and antimicrobial susceptibility test (AST). Demographics such as sex, age and other patients’ information were extracted from patient request forms. Patients’ clinical outcomes were retrieved from patient’s clinical case notes.

### Blood culture and bacterial identification

Blood for culture were collected by attending clinician into blood culture bottles for adult (BD BACTEC Plus Aerobic /F Culture Vials, Becton Dickinson and Company) and pediatric (BD BACTEC Peds Plus^™^/F Culture Vials, Becton Dickinson and Company). Upon reaching the laboratory, blood culture bottles were inspected for acceptance criteria. Blood culture vials were incubated into BD BACTEC FX40 analyzer for maximum of five days.

Primary Gram stain was performed on positive cultures followed by subculture culture on appropriate solid culture media. A single drop of blood was inoculated into 5 % sheep Blood Agar and MacConkey agar (MCA), then incubated at 37°C with 5-10% CO_2_ and 37°C respectively for duration of 18-24 hours. Bacteria were initially identified by colonial morphology and Gram stain. Gram-positive cocci were further identified by a set of biochemical tests including, catalase test, coagulase, DNase, Staphaurex, *Streptococcus* Grouping kit. Gram negative rods were further identified by API20 E and API20 NE (Biomerieux, France).

### Antimicrobial susceptibility testing

Kirby Bauer disc diffusion method was used to test antimicrobial susceptibility following Clinical and Laboratory Institute guidelines [CLSI] (9). Antibiotics included in susceptibility testing were those commonly used in our settings for management of BSI and few reserved for severe bacterial infections.

Multidrug resistance (MDR) was defined as resistance to at least one antibiotic in three or more antimicrobial classes (10). MRSA was determined by disc diffusion method using cefoxitin disk (30μg) as previously described (9). ESBL production in Enterobacteriaceae was detected as described previously (9). Carbapenemase production in Gram negative bacteria was screened using combination disk method where by meropenem disk (10μg) alone and combination of meropenem disk (10μg) with dipicolinic acid (DPA) 1000μg. An increase ≥5 mm in zone diameter around combined meropenem disk and DPA, as compared with the disk of meropenem alone, was considered to be a positive result (11).

The following reference strains were used for quality control: *Escherichia coli* ATCC 25922, *K. pneumoniae* ATCC 700603 for ESBL, *S. aureus* ATCC 25923, and *S. aureus* ATCC 29213 for MRSA and *K. pneumoniae* ATCC 1705 and *K. pneumoniae* ATCC 1706 for Carbapenamase resistance

### Data analysis

Descriptive analysis for categorical variables was summarized in form of frequencies and percentages. The comparison within variables was performed using Chi-square test or Fisher’s exact test to observe significances of proportion differences. Binary univariate regression analysis was performed to identify factors associated with mortality. All variables with *p* < 0.2 at univariate analysis were further analyzed in multivariate binary regression to identify independent factors associated with mortality. P value < 0.05 was considered statistically significant.

### Ethical approval

Ethical approval for the study was obtained from Senate Research and Publication committee, the Institutional Review Board of Muhimbili University of Health and Allied Science. Permission to conduct study at Muhimbili National Hospital was granted by MNH executive director.

## Results

### Description of the study participants

A total of 402 specimens for blood cultures were included in the study. Majority (39.3%) were obtained from patients aging 1-14 years and 235 (58.5%) were from males. Most blood cultures (40.8%) were from pediatric wards while 16.9% (68/402) were from patients admitted in ICU. A total of 87(21.6%) participated died during in the study period (Table 1).

**Table 1:** Demographic and Clinical characteristics of study participants

| Variable | Frequency | Percentage |
| --- | --- | --- |
| **Age** |  |  |
| <1 month | 53 | 13.2 |
| 1-14 years | 158 | 39.3 |
| 15-46 | 113 | 28.1 |
| >46 | 78 | 19.4 |
| **Sex** |  |  |
| Male | 235 | 58.5 |
| Female | 167 | 41.5 |
| **Ward** |  |  |
| Pediatric | 164 | 40.8 |
| Surgical | 79 | 19.7 |
| Medical | 91 | 22.6 |
| ICU | 68 | 16.9 |
| **Outcome** |  |  |
| Death | 87 | 21.6 |
| Discharged | 315 | 78.1 |

### Rate of Laboratory confirmed blood-stream infections

Of all 402 participants with blood culture included in the study, 11.4% (46/402) (95%CI 8.6 – 15) had culture positive BSI, and 17 of these patients died (case fatality rate was 37%). The rate of culture positive BSI were more in female (12%, 20/167) than in male (11.1%, (26/235). There was no significant difference in rate of BSI in neonate (15.1%, 8/53), children aged 1-14 years (7.6%, 12/158), patients aged 15 – 46 years (13.3%, 15/113) and patients aged more than 46 years (14.1%, 11/78), p=0.3. Patients admitted at ICU had high rate of BSI (17.6%, 12/68) compared to those admitted at pediatric (8.5%, 14/164), surgical (8.9%, 7/79) and medical ward (14.3%, 13/91) p=0.2. There was a significant high rate of culture positive BSI in deceased patients (19.5%) compared to those survived (9.2%), p =0.008.

### Bacterial isolates and antimicrobial susceptibility pattern

A total of 46 pathogens were isolated from blood cultures. Majority (74%) were Gram-negative bacteria, of which *K. pneumoniae* and *P. aureginosa* were the commonest accounting for 23.9% each. The most predominant pathogen among the Gram-positives was *Staphylococcus aureus* (10/12) (figure 1). Antimicrobial susceptibility test was performed for 43 bacterial isolates to determine susceptibility pattern. Majority of isolates (70.5%, 31/44) were multi-drug resistant. Enterobacteriaceae displayed high rates of resistance to multiple antibiotics tested as presented in table 2.

**Figure 1:**
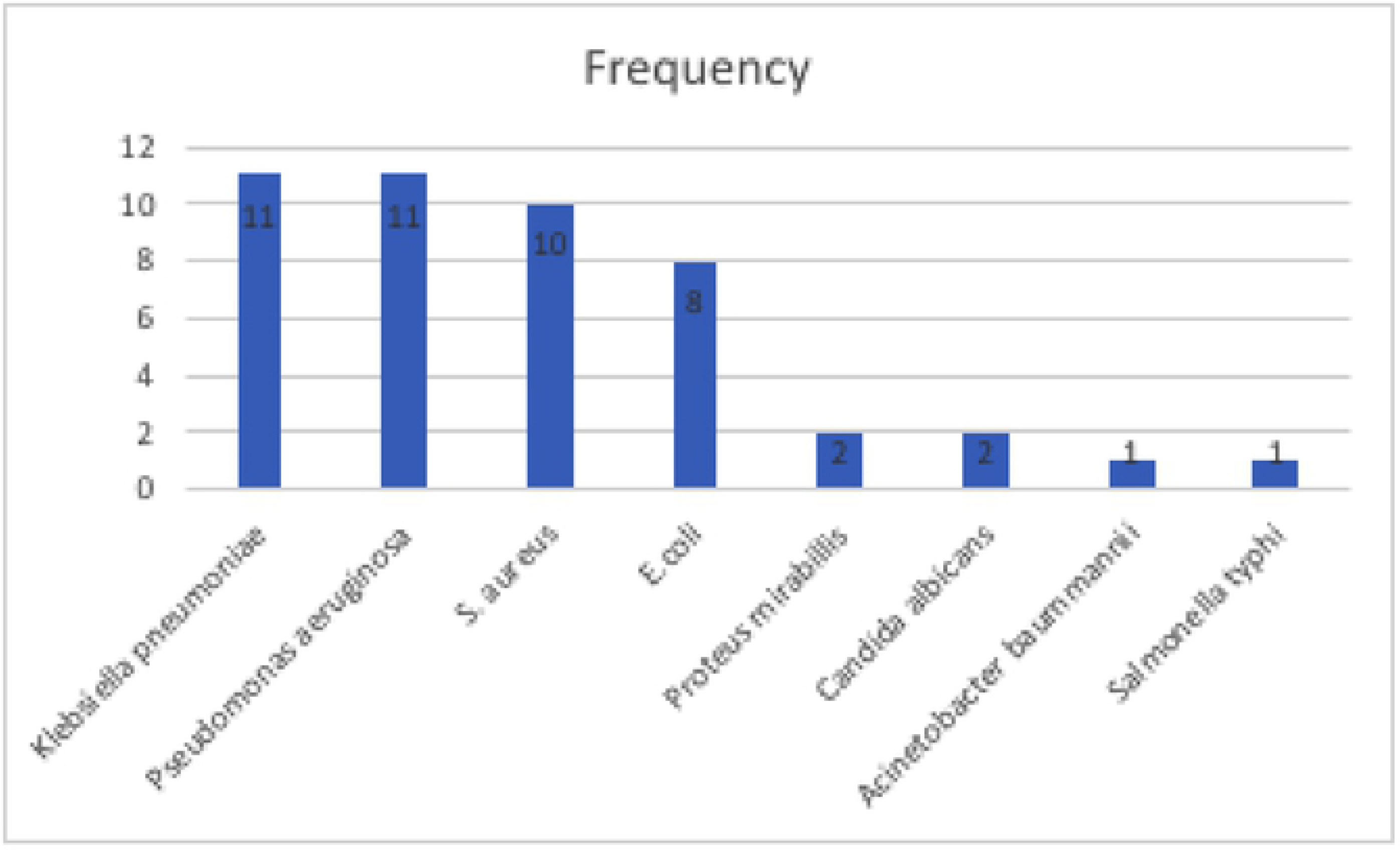
Bacteria isolated from patients with BSI at MNH.

**Table 2:** Antimicrobial resistance pattern of bacteria isolated from patients with BSI at MNH

| Bacteria species | N | Percent of bacteria resisted |  |  |  |  |  |  |  |  |  |  |  |  |  |  |
| --- | --- | --- | --- | --- | --- | --- | --- | --- | --- | --- | --- | --- | --- | --- | --- | --- |
|  |  | CN | CR<br>O | CAZ | CTX | AM<br>C | AK | MEM | PR<br>T | CIP | SXT | FOX | DA | P | DO<br>X | E |
| *E. coli* | 8 | 25 | 63 | 57 | 88 | 71 | - | - | 25 | 50 | 50 | - | - | - | - | - |
| *K. pneumoniae* | 11 | 55 | 89 | 90 | 80 | 91 | 9 | - | 20 | 36 | 80 | - | - | - | - | - |
| *Proteus mirabilis* | 2 | 100 | 100 | 100 | 100 | 100 | 100 | 100 | - | 100 | 100 | - | - | - | - | -- |
| *P. aeruginosa* | 11 | 73 | 100 | 70 | 100 | - | 55 | 46 | 46 | 64 | - | - | - | - | - | - |
| *S. aureus* | 10 | 44 | - | - | - | - | - | - | - | 70 | 10 | 40 | 20 | 80 | 60 | 70 |
| *Acinetobacter* species | 1 | - | 100 | 100 | 100 | 100 | - | 100 | 100 | 100 | - | - | - | - | - | - |

Key: CN gentamicin, CRO ceftriaxone, CAZ ceftazidime, CTX cefotaxime, AMC Amoxicillin-clavulanic acid, AK amikacin, MEM meropenem, PRT Piperacillin-tazobactam, CIP ciprofloxacin, SXT trimethoprim-sulfamethoxazole, DA clindamycin, P penicillin, DOX doxycycline, E erythromycin

Almost Sixty eight percent (68.2%, 15/22) of Enterobacteriaceae were ESBL producers. *Pseudomonas aeruginosa* showed over 60% to 100% of resistance to commonly prescribed antibiotics including: gentamicin (73%), ceftriaxone (100%), cefotaxime (100%), ceftazidime (70%) and ciprofloxacin (64%). Remarkably, 46% of *Pseudomonas aeruginosa* were resistance to meropenem; furthermore, 27% (3/11) of them were Carbapenemase producing pathogens. Resistance to Piperacillin-tazobactam, uncommonly used antibiotic were observed in 25% of *E. coli*, 20% of *Klebsiella pneumoniae* as well as 46% of *Pseudomonas aeruginosa. Staphylococcus aureus* displayed high resistance to ciprofloxacin (70%), Penicillin (80%) and erythromycin (70%); in addition, (40%) of *S. aureus* were MRSA.

### Predictors of mortality in blood stream infection

On univariate analysis, patients with positive blood culture were two times more likely to die compared to those with negative blood culture (cOR 2.39, 95%CI 1.24 – 4.59, *p* =0.009). Patients aged more than 14 years suspected of BSI were at more odds of dying compared to those below (cOR 2.26, 95% 1.38 – 3.69, *p* = 0.001). Patients suspected of BSI admitted at surgical ward (cOR 2.61, 95%CI 1.32 – 5.18, *p* = 0.006) and ICU (cOR 2.57, 95%CI 2.57 – 9.87, *p* < 0.001) were at high risk of dying compared to those admitted at medical ward. BSI with multi-drug resistant bacteria were found to predict mortality (cOR 6.1, 95%CI 1.2 – 31.6, *p* =0.03) than non-MDR infection (table 3).

**Table 3:** Predictors of Mortality in Patients with suspected and confirmed Blood stream infection

| Variable | Frequency | Death %(n) | cOR | 95%CI | p value | aOR | 95%CI | p value |
| --- | --- | --- | --- | --- | --- | --- | --- | --- |
| **All patients with suspected BSI (n=402)** |  |  |  |  |  |  |  |  |
| **Blood culture** |  |  |  |  |  |  |  |  |
| Negative | 356 | 19.7(70) | 1 |  |  | 1 |  |  |
| Positive | 46 | 37(17) | 2.39 | 1.24 – 4.59 | 0.009* | 2.18 | 1.1 – 4.31 | 0.03* |
| **Sex** |  |  |  |  |  |  |  |  |
| Female | 117 | 19.9(33) | 1 |  |  |  |  |  |
| Male | 235 | 23(54) | 1.2 | 0.74 – 1.97 | 0.44 |  |  |  |
| **Age (years)** |  |  |  |  |  |  |  |  |
| <14 | 211 | 15.2(332) | 1 |  |  | 1 |  |  |
| >14 | 191 | 28.8(55) | 2.26 | 1.38 – 3.69 | 0.001* | 0.8 | 0.39 – 1.62 | 0.5 |
| **Ward** |  |  |  |  |  |  |  |  |
| Pediatric | 164 | 12.2(20) | 1 |  |  |  |  |  |
| Surgical | 79 | 26.6(21) | 2.61 | 1.32 – 5.18 | 0.006* | 2.16 | 0.87 – 5.36 | 0.1 |
| Medical | 91 | 19.8(18) | 1.78 | 0.89 – 3.56 | 0.12 | 1.43 | 0.6 – 3.43 | 0.4 |
| ICU | 68 | 41.2(28) | 5.04 | 2.57 - 9.87 | <0.001* | 4 | 1.7 – 9.41 | 0.001* |
| **Patients with BSI (n=46)** |  |  |  |  |  |  |  |  |
| **MDR** |  |  |  |  |  |  |  |  |
| No | 15 | 13.7(2) | 1 |  |  | 1 |  |  |
| Yes | 31 | 48.4(15) | 6.1 | 1.2 – 31.6 | 0.03* | 7.4 | 1.24 – 43.83 | 0.03* |
| **ESBL** |  |  |  |  |  |  |  |  |
| No | 31 | 35.5(11) | 1 |  |  |  |  |  |
| Yes | 15 | 40(6) | 1.2 | 0.34 – 4.31 | 0.7 |  |  |  |
| **Organism type** |  |  |  |  |  |  |  |  |
| GNR | 36 | 35.3(12) | 1 |  |  |  |  |  |
| GPC | 12 | 41.7(5) | 1.31 | 0.34 – 5.03 | 0.7 |  |  |  |

cOR = crude odds ratio, aOR = adjustable odd ration, 95%CI= 95% Confidence interval, \* significant

On multivariate analysis, result of blood culture, location of admission and status MDR were found to be independent risk factors for mortality in patients suspected of BSI. Positive bacteria blood culture, had 2 times more odds of mortality than blood culture negative (aOR 2.18, 95%CI 1.1 – 4.31, p 0.03). Admission at ICU remained independent risk factor for mortality with 4 times odds of dying compared with admission at pediatric wards (aOR 4, 95%CI 1.7 – 9.41, p 0.001). Presence of Multi-drug resistant bacteria was found to be independent predictor for mortality with 7 times odds of dying compared to non MDR bacterial infection (aOR 7.4, 95%CI 1.24 – 43.83, p 0.03) Table 3.

## Discussion

In this study conducted at tertiary hospital setting, we found slightly lower (11.4%) prevalence of BSI compared to previous findings at the same facility (1, 12). However, our study observed significantly high case fatality rate of 37%. Reports from various study populations have reported different prevalence rates of BSI (1, 4, 5, 12–16). Blomberg et al. found the prevalence of 13.9% among admitted children at the same hospital (1). Likewise, Moyo et al. reported the prevalence of 13.4% among all patients of different age groups at the same hospital (12). Moyo et al included Coagulase negative *Staphylococcus* as the true pathogens in the analysis, which were not considered in our study. All patients included in the study had used antibiotics prior to blood culture, which might explain the observed low rate of BSI. In this study, all patients had hospital associated BSI; in this regard, the hospital need to intensify infection prevention and control program to limit the spread of BSI and its associated mortality.

As expected, patients admitted in ICU suspected with BSI had high risk of mortality compared to those admitted in non-ICU wards. This finding is in line with other earlier studies on risk factors for BSI (6, 8). We observed that unlike patients admitted in non-ICU, patients in ICU were likely to have other underlying diseases. This observation could explain the observed high mortality. In these circumstances, our findings suggest prompt investigation in suspected BSI in ICU and appropriate antibiotic use guided by laboratory results.

Our study observed that positive bacterial blood culture was an independent laboratory predictor for mortality in patients suspected of BSI. Similar finding were also observed in previous studies from developing countries in neonate (17), children (13), and adults (2). This finding emphasizes the important of blood culture in suspected BSI. It also highlights the role of laboratory in prompt notifying clinician when blood culture is positive as it could guide early choices of empiric antibiotics for better treatment outcomes. In addition, when pathogen is identified and susceptibility results are available, clinician need to be alerted for them to adjust/de-escalation of antibiotics. With overall BSI case fatality rate of 37%, further study needs to be performed to evaluate the impact of laboratory prompt notification of results. Consideration of duration of blood culture to positivity on antimicrobial stewardship can have impact on patients’ outcomes.

Infection due to multi-drug resistance bacteria was an independent predictor of mortality in our study. This finding was comparable to previous studies in Africa and elsewhere (2, 18). Treatment of these infections is very difficult and carries poor prognosis as bacteria are resistant to all available antibiotics’ options. Patients infected with MDR pathogens were supposed to receive reserve antibiotics like vancomycin, carbapenems or colistin; unfortunately, they are expensive and unavailable in our settings. Although performing and reporting phenotypic AST and defining MDR pathogens remain crucial for patients care, it is not often done in our settings. Our finding justifies the need for clinical microbiology laboratory to notify and discuss with clinician whenever they isolate MDR bacterium. In addition, clinical microbiologists and infectious diseases specialists need to work hand in hand in the management of MDR infected patients.

The current study observed that bacteria causing BSI were highly resistant to most antibiotics commonly used in our setting. The trend of resistance were compatible to resistance patterns observed in previous studies at the same setting (1, 12) and other hospital associated infections (19). Persistent of high rates of resistance at our setting could be accounted by increasingly empiric use of antibiotics; in most cases imprudent use of antibiotics. Lately, clinical microbiology laboratory does not produce annual AMR report, which could have guided clinician in empiric prescription pattern. Understanding of local epidemiological AMR patterns is critical for informed prescription practice. We strongly recommend the clinical microbiology laboratory to regularly produce local AMR reports.

The study revealed 68.2% of Enterobacteriaceae being phenotypic confirmed ESBL producers. Blomberg et al. reported high rates of genotypic confirmed ESBL producing Enterobacteriaceae in children with BSI at the same hospital that predicted mortality (3). Similarly, high rates of ESBL producing pathogens has been reported at the same hospital from clinical isolates of urine (20) and surgical site infections (19). Beside ESBL, we also observed MRSA and phenotypic carbapenemase production in *Pseudomonas aeruginosa* and *Proteus mirabilis*. Carbapenemase producing *Pseudomonas aeruginosa* have also been reported previously at our hospital (21). It is worrying to note the growing resistance to carbapenems, which is the last effective therapy for severe Gram-negative infections. As observed in earlier studies, ESBL, MRSA and carbapenemase producing isolates carries a very poor prognosis and associated with increased health care cost (3, 17). The findings warrant the need for clinical microbiology laboratory to enforce policy of detecting and reporting resistance pathogens. If enforced, will help in early detection of an outbreak due these pathogens.

### Conclusion

The overall prevalence of BSI was 11.4%, and 17 of patients died (case fatality rate was 37%). Majority of the bacteria isolated from BSI were multi-drug resistant. Admission to ICU and positive blood culture were independently associated with mortality in suspected BSI. Multi-drug resistance bacteria were an independent predictor for mortality in confirmed BSI.

## Acknowledgment

Authors would like to thank the management of Muhimbili National Hospital for the support on reagents, supplies and other consumables which were used in the study.

## Authors contribution

JM conceived the study, performed analysis and drafted the manuscript, UK edited the manuscript, EM collected data and performed experiment and MM reviewed and edited the manuscript.

